# Tetraspanin positivity as a function of extracellular vesicle size measured by a modified immuno-TEM protocol

**DOI:** 10.1101/2025.03.13.643132

**Authors:** Pablo Fagúndez, Álvaro Olivera, Olesia Gololobova, Marco Li Calzi, Alfonso Cayota, Kenneth Witwer, Eduardo Méndez, Juan Pablo Tosar

**Affiliations:** Unidad de Bioquímica Analítica, Centro de Investigaciones Nucleares, Facultad de Ciencias, Universidad de la República, Uruguay; Laboratorio de Biomateriales, Instituto de Química Biológica, Facultad de Ciencias, Universidad de la República, Uruguay; Functional Genomics Laboratory, Institut Pasteur de Montevideo, Uruguay; PEDEClBA - Program for the Development of Basic Sciences.; Centro Universitario Regional Este, Universidad de la República, Uruguay; Department of Molecular and Comparative Pathobiology, Johns Hopkins University School of Medicine, Baltimore, Maryland, USA; Hospital de Clínicas, Universidad de la República, Uruguay; Department of Neurology, Johns Hopkins University School of Medicine, Baltimore, Maryland, USA; The Richman Family Precision Medicine Center of Excellence in Alzheimer’s Disease, Johns Hopkins University School of Medicine, Johns Hopkins Medicine and Johns Hopkins Bayview Medical Center, Baltimore, Maryland, USA

## Abstract

The identification of surface markers that correlate with specific subpopulations of extracellular vesicles (EVs) is important for EV identification, classification, purification, sorting, and functional analysis. Tetraspanins such as CD9, CD63 and CD81 were once considered to be universal markers of exosomes: small EVs released into the extracellular space when late endosomes / multivesicular bodies fuse with the plasma membrane. In contrast, plasma membrane-derived ectosomes (also called microvesicles) have a different biogenesis, were often regarded as being larger than exosomes, and display a different surface proteome. However, recent studies have shown that tetraspanins such as CD9 and CD81 are highly enriched on ectosomes derived from various sources. Thus, it is currently unclear how tetraspanin content correlates with specific EV subpopulations. Here, we present a modified immuno-TEM protocol that can be easily applied to heterogeneous EV populations comprising both small and large EVs (and presumably also a collection of exosomes and ectosomes). In EVs purified from U-2 OS cells by size-exclusion chromatography, we show that the percentage of particles positive for CD9 and CD81 is significantly higher in the subpopulation of EVs ≤ 100 nm (i.e., small EVs). These results also explain discrepancies in the size distribution profiles that we obtained using the same EV preparations by alternative single-vesicle characterization platforms such as nano flow cytometry and SP-IRIS / ExoView. The latter, when used to capture tetraspanin-positive particles, returns a population that is relatively small in size.

## 1. Introduction

Extracellular vesicles (EVs) are cell-derived nanoparticles delimited by a lipid bilayer. They fulfill several important roles in biological systems, including disposal of cellular waste and as mediators of intercellular communication pathways (Couch et al., 2021). While not strictly defined by the presence of specific surface markers, EVs frequently display tetraspanins such as CD9, CD81, and CD63, in addition to various cell-specific and -nonspecific surface and intraluminal proteins, RNA, and other macromolecules (Jeppesen et al., 2019; Kowal et al., 2016; Mathieu et al., 2021).

EVs derived from non-apoptotic cells can be classified based on their biogenesis as either exosomes or ectosomes. Exosomes are derived from endocytic compartments while ectosomes (sometimes referred to as microvesicles) are derived by direct budding from the plasma membrane (Welsh et al., 2024). EVs can also be classified according to size, e.g., as small EVs and large EVs, a rough and often arbitrary classification that initially arose from the assumption that EVs are easily separated by size using differential ultracentrifugation. An important source of confusion in the field is the widespread assumption that small EVs and large EVs tend to equate with exosomes and ectosomes, respectively (Keerthikumar et al., 2015). However, this is not necessarily the case. While some ectosomes are larger than exosomes (which are limited by the size of multivesicular bodies), the two EV types overlap completely in the 30 – 150 nm range. Therefore, exosomes and small ectosomes co-purify in small EV preparations (Cocucci & Meldolesi, 2015; Jeppesen et al., 2023; Mathieu et al., 2021)

This confusion, together with an inconsistent use of nomenclature across studies, is further complicated when it comes to defining molecular markers specific to exosomes/ectosomes and small/large EVs. For instance, the tetraspanins CD9, CD63 and CD81 have often been considered markers of “classical exosomes” (Jeppesen et al., 2019), and tend to be more enriched in small EVs collected by differential ultracentrifugation (Kowal et al., 2016). However, at least for HeLa cells as an EV source, CD9 and CD81 tend to be more associated with small ectosomes derived from the plasma membrane, while CD63 is more enriched in *bona fide* exosomes (Mathieu et al., 2021). Even so, no tetraspanin may be 100% specific for either plasma membrane or endosomal membrane.

If exosomes and ectosomes cannot be segregated by either their physicochemical or their biochemical characteristics, operational terms like small EVs and large EVs become more convenient when working with extracellular samples. However, it is still not clear whether there is a strong correlation between EV size and tetraspanin positivity. As mentioned earlier, tetraspanins tend to be more enriched in 100,000 x g pellets (i.e., presumed small EV) compared with 10,000 x g pellets (i.e., presumed large EVs), but CD63 and CD9 are also strongly detected in 2,000 x g pellets, corresponding to very large EVs (Kowal et al., 2016). In addition, the presence of CD9 and CD81 in small ectosomes derived from the plasma membrane (Mathieu et al., 2021) suggests that they could also be present in larger ectosomes released by a similar mechanism.

If there is a size cut-off that can distinguish EVs enriched or not with tetraspanins, what that hypothetical size cut-off be? This question cannot be answered by simply comparing between 10K and 100K x g pellets, because this method is too coarse-grained if different EVs subpopulations present broad size distributions, as seems to be the case. Here, we developed an unbiased determination of CD9 and CD81-positivity by a modified immuno-TEM protocol, using EVs purified by size exclusion chromatography (SEC) without imposing arbitrary size cut-offs by prior centrifugation steps. Our results clearly show that these two tetraspanins are strongly enriched in EVs < 100 nm, which could correspond to small exosomes, ectosomes, or both.

## 2. Results

### 2.1 Cross-platform EV sizing shows diminished detection of large EVs by SP-IRIS

Nanoparticle sizing platforms can have a profound impact on measured EV size distributions (Arab et al. 2021). To obtain a platform-independent “ground-truth”, we used SEC to purify EVs from the conditioned medium of U-2 OS cells and sized more than 300 individual particles with EV-consistent morphology by high resolution TEM (**Figure 1A**, inset). Unlike several publications wherein the cell-conditioned media (CCM) was centrifuged at 10,000 – 100,000 x g before separation by SEC, we concentrated the CCM by ultrafiltration and loaded it directly into the columns. As a result, our EVs preparations should contain a mixture of large and small EVs in the same proportions in which they were present in the CCM (except for any EVs lost during the 2,000 x g spin).

**Figure 1:**
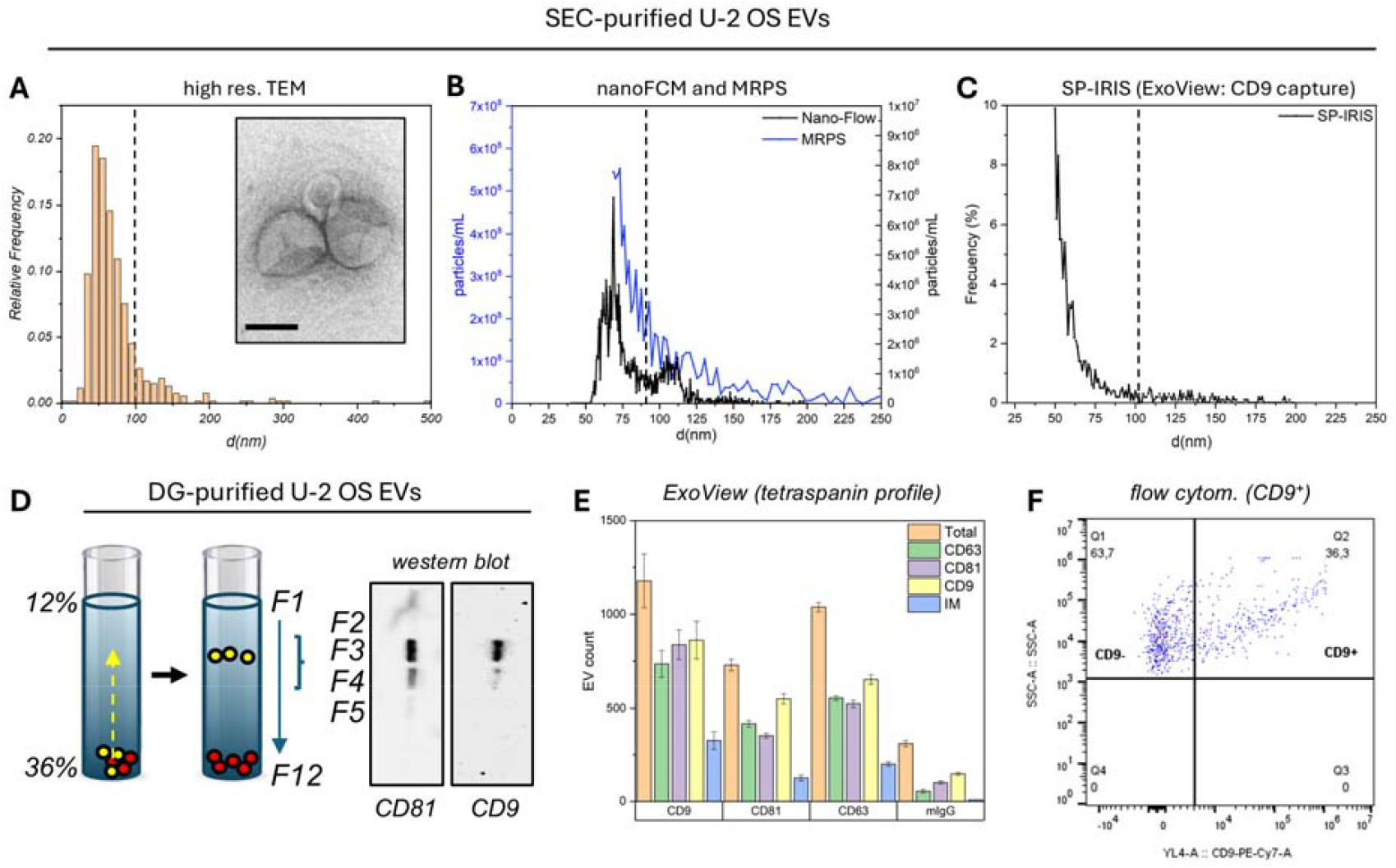
Characterization of SEC-purified U-2 OS EVs. The size distribution of EVs was determined by high-resolution TEM (A), MRPS and nanoFCM (B) and SP-IRIS (C). The inset in (A) shows a representative micrograph. EVs were also purified by density, using iodixanol flotation gradients, and analyzed by western blot (D). The levels of the tetraspanins CD9, CD81, and CD63 were determined by ExoView (E). The X axis separates categories based on the capture antibody, while the different colors represent the results based on fluorophore-labeled detection antibodies. The percentage of CD9-positive EVs was also determined by conventional flow cytometry (F). All EVs in this figure were purified by SEC, except for those in panel (D).

EV sizing by TEM followed a bimodal distribution, with most particles ranging from 30 to120 nm (mode at: 50 - 60 nm), and a second population of larger EVs ranging from 100 to 170 nm (**Figure 1A** and **Figure S1, A**). Particles larger than 170 nm in diameter were observed but were infrequent. Thus, while both small (≤ 100 nm) and large EVs (> 100 nm) are present in U-2 OS CCM, the former population is much more abundant. Of note, a mode of 55 nm was identified for DiFi cell-derived EVs using a single-molecule flow cytometer with a limit of sizing ≤ 35 nm (Kim et al., 2024). We are therefore confident that the size distribution obtained here by TEM reasonably reflects the real size distribution of the particles.

Nano flow cytometry (nanoFCM) showed a size distribution consistent with TEM results (**Figure 1B**), with a major peak at 60 – 70 nm and a smaller peak at 100 – 120 nm. The ratio between the two peaks was smaller than by TEM, but still similar. In contrast, the distribution returned by microfluidic resistive pulse sensing (MRPS) started at 70 nm due to our inability to distinguish smaller particles from background but reproduced the “tail” of the TEM distribution. While no distinct peak corresponding to large EVs was evident, a substantial number of events with a high signal-to-noise ratio were > 100 nm (**Figure 1B**, blue line).

Surprisingly, SP-IRIS showed a distribution decaying from 50 nm onwards, with virtually no particles > 100 nm (**Figure 1C**). The absence of the large EV population was unexpected, considering that larger particles should be easier to detect by interferometric imaging. However, SP-IRIS, as implemented by the capture chips we used with the ExoView platform, captures EVs using anti-tetraspanin antibodies. Size-distribution profiles were similar for EVs captured by CD9 (**Figure 1C**), CD63, and CD81 (**Figure S1, B**). We therefore considered the possibility that surface-exposed tetraspanins are not distributed evenly between EVs of different sizes, with the smallest EVs being more enriched in these proteins.

### 2.2 Detection of CD9 and CD81 on the surface of SEC-purified U-2 OS EVs

We have previously isolated U-2 OS EVs by isopycnic centrifugation (Tosar et al. 2020) and showed the presence of CD9 and CD81 in these EVs by Western blot (**Figure 1D**). The ExoView platform also confirmed the presence of CD9, CD63, and CD81 on the surface of SEC-purified EVs, with CD9 > CD63 ≈ CD81 (**Figure 1E**), and with most particles being triple-positive (**Figure S1, C-D**).

Although the ExoView platform allows us to determine the relative proportions of each tetraspanin on the surface of individual EVs, tetraspanin-negative EVs are not captured, potentially leaving an important population of EVs unmesured. To circumvent this problem, we labeled EVs with a PE-Cy7-conjugated anti-(h)CD9 antibody and analyzed particles by flow cytometry. Here, we found that approximately 36% of gated particles (based on side-scatter, SSC, using a 488 nm laser) were positive for CD9 (**Figure 1F** and **Figure S2**). All gated events disappeared when pre-treating the samples with detergent (**Figure S2, A**), suggesting that the measured events were indeed lipid-containing vesicles. Unfortunately, the fluorescence intensity of anti-CD81-labeled EVs was too dim to reliably measure these particles (data not shown). In addition, a major caveat of the flow cytometry instrument that we used for this determination (Attune NxT) is that it can detect only particles that are larger than 100 nm in diameter. Thus, the CD9 positivity rate calculated with this instrument does not correspond to the entire population of particles, because small particles are below the detection limit of the instrument (**Figure 1A-C**).

### 2.3 Synthesis and characterization of AuNP-AG as a reagent for immuno-TEM

To determine the positivity rate of CD9 and CD81 in the most unbiased way possible, and to correlate labelling rates with particle size, we performed immuno-TEM with SEC-purified EVs. To do this, we first synthesized 13 nm gold nanoparticles (AuNP-cit) and modified them with protein A/G to capture anti-tetraspanin antibodies. The immobilization of protein A/G on the surface of freshly prepared AuNPs followed the step-by-step quality-control guide that we presented earlier (Fagúndez et al. 2021).

The synthesized AuNP-cit had an average diameter of 13 ± 4 nm as determined by TEM (**Figure 2A**). The plasmon absorption peak was observed at 522.2 ± 0.2 nm, characteristic of AuNPs of this size (**Figure 2B**). Dynamic light scattering (DLS) showed a single, monodisperse population (pdi = 0.273) with an average diameter of 16 ± 5 nm (**Figure S3, A**).

**Figure 2:**
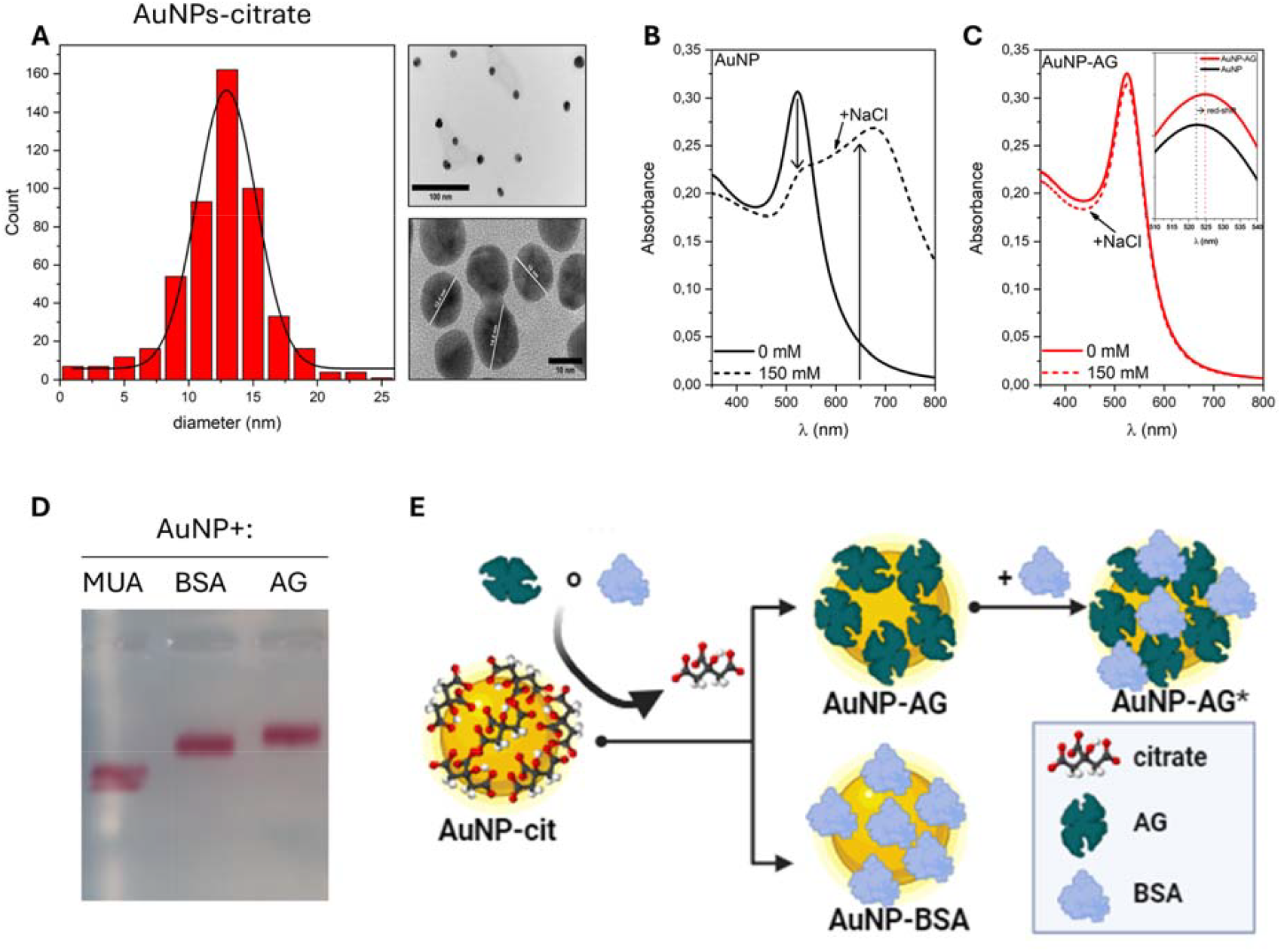
Synthesis and characterization of protein-modified gold nanoparticles (AuNPs). The size distribution and a representative micrograph of the synthesized AuNPs are shown (A). The distribution was fitted to a normal distribution. The UV-visible spectra of citrate-coated (B) or protein A/G-coated (C) AuNPs, before and after NaCl addition, is also shown. The inset shows the 3 nm shift in the absorption maximum, expected for protein-modified AuNPs. The electrophoretic mobility for AuNP-AG and AuNP-BSA is shown in (D). The delay in migration compared to non-protein-modified AuNPs (AuNP-MUA) is noticeable. Panel (E) depicts the different, consecutive stages for the obtention of either AuNP-BSA (negative control reagent) and AuNP-AG, with BSA blocking available sites in the gold nanoparticle (AuNP-AG*).

A hallmark of unmodified AuNP-cit is their tendency to aggregate at high ionic strengths, changing color from red to blue (Fagúndez et al. 2021). This can be observed as a sharp increase in absorbance at 650 nm and a decrease at 520 nm when incubated in 150 mM NaCl (**Figure 2B**). However, incubation for 24 h with either protein A/G or BSA rendered the modified nanoparticles completely insensitive to salt-induced aggregation (**Figure 2C**). In addition, a small red-shift of the plasmon maxima was observed, what is often considered indicative of successful ligand exchange. DLS confirmed an expected increase in the hydrodynamic diameter for both AuNP-AG and AuNP-BSA (55 ± 10 nm and 25 ± 8 nm, respectively; **Figure S3, B**).

Electrophoretic mobility was also decreased in protein-modified AuNPs compared to those modified with a smaller ligand (MUA), consistent with successful protein immobilization (**Figure 2D**). Note that we could not use citrate-modified AuNPs because they immediately aggregate when in contact with the electrophoresis buffer (Fagúndez et al. 2021). To block potential protein-free sites in AuNP-AG, nanoparticles were centrifuged to remove excess protein A/G and resuspended in PBS-BSA for up to 1 h (**Figure 2E**; AuNP-AG*). Centrifuged and resuspended nanoparticles were slightly more retarded in gel electrophoresis than their non-centrifuged counterparts (**Figure S4**). However, they still migrated as discrete red bands (rather than a smear, which would indicate aggregation). In addition, DLS analysis did not suggest centrifugation-induced aggregation (data not shown).

### 2.4 A modified immuno-TEM protocol for EV phenotyping

Standard immuno-TEM assays employ AuNPs conjugated to antibodies, or to proteins like streptavidin, capable of incorporating antibodies *a-la-carte*. However, the ∼13 nm AuNP-cit obtained with our synthesis method were found to be unstable in most immunoassay buffers once modified with streptavidin (Fagúndez et al. 2021). Conjugation to IgG rendered relatively stable nanoparticles, but these aggregated upon the removal of free ligands by centrifugation (**Figure S4**). In contrast, AuNPs modified with protein A/G were stable even in high ionic strength conditions (**Figure 2C**) and could be centrifuged and resuspended in PBS without substantial aggregate formation (**Figure S4, A**). Unfortunately, incubation of AuNP-A/G with IgGs destabilized the nanoparticles after the centrifugation step (**Figure S4, B**), despite our use of various resuspension conditions and buffers.

To solve the AuNP instability problem, we designed a different protocol, in which EVs were first incubated with anti-CD9 or anti-CD81 antibodies, purified by SEC, and applied onto TEM grids (**Figure 3A**). The grids were then blocked with BSA, and AuNP-AG* were directly applied to the grids containing antibody-labeled EVs (**Figure 3B**). AuNP-BSA and EVs labeled with an irrelevant antibody (anti-C23, isotype control; **Figure 3C**) served as negative controls to evaluate unspecific binding and optimize blocking conditions (**Figure S5)**, washing steps, incubation times, and nanoparticle dilutions (see methods).

**Figure 3:**
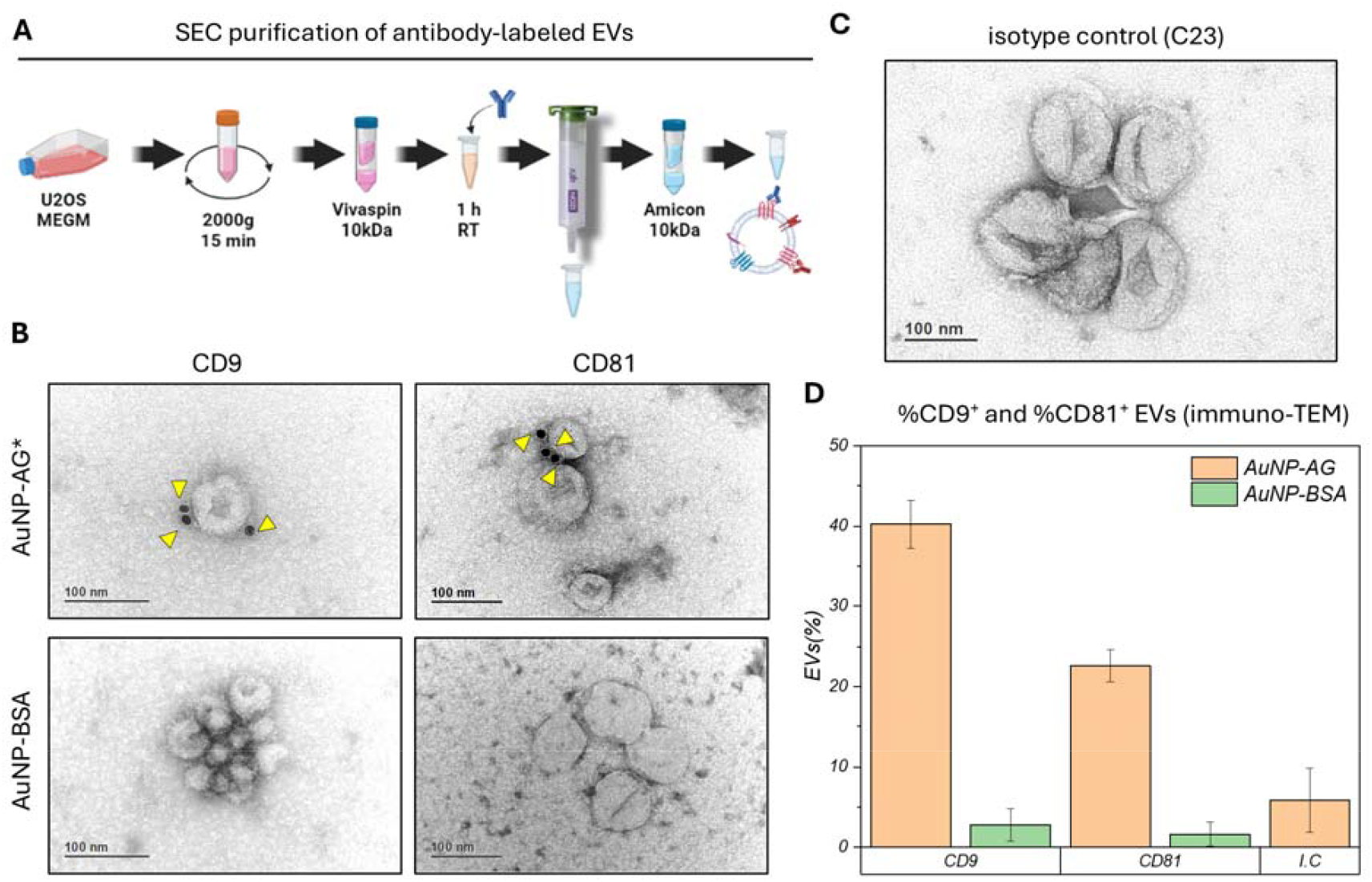
Detection of CD9 and CD81 by our modified immuno-TEM protocol. Experimental procedure followed for labeling and purification of EVs prior to immuno-TEM (A). TEM micrographs of EVs labeled with either an anti-CD9 or an anti-CD81 antibody (B), or an anti-C23 isotype control (C), and treated with either AuNP-AG* (B, C) or AuNP-BSA (B) on a TEM grid. The yellow triangles indicate the presence of AuNPs on the surface of EVs. The percentage of EVs bound to AuNPs for the different antibodies and AuNPs were calculated (D). Error bars correspond to two independent replicates done on different days with different batches of EVs and AuNPs.

Quantitative analysis of N = 300 EVs (with and without AuNPs) showed that approximately 40% of EVs treated with anti-CD9 were positive for AuNP-AG*, while this figure decreased to 22% for those treated with anti-CD81 (**Figure 3D**). This was virtually identical to the numbers obtained by flow cytometry (36% for CD9; **Figure 1F**), despite this technique being biased against small particles. The fact that CD9 > CD81 was also consistent with ExoView results (**Figure 1E**). In parallel, negative controls (AuNP-BSA and AuNP-AG* with EVs labeled using an isotype control) were all below 6% (**Figure 3, D**).

In addition to determining tetraspanin positivity, we counted the number of AuNPs attached to each EV. Interestingly, while most CD81-labeled EVs had a single AuNP-AG*, there was a strong tendency to more frequently record EVs with 2 – 5 AuNPs when the EVs were labeled with the anti-CD9 antibody (**Figure 3B** and **Figure S6, A**). Thus, CD9 is not only more frequently detected on U-2 OS EVs than CD81; it is also present in higher copy numbers per vesicle (assuming similar antibody binding efficiency). This could explain, for example, why we failed to reliably detect CD81 by flow cytometry despite robust detection of CD9^+^ particles.

### 2.5 Tetraspanin positivity as a function of EV size

A representative micrograph of an EV cluster comprising particles of different sizes shows that AuNP-AG* binding predominantly occurs in EVs ≤ 100 nm (**Figure 4A** and **Figure S6, B**). The left image (i.e., anti-CD9-labeled EVs) depicts 4 large EVs and 12 small EVs. None of the former are associated with AuNPs, while 6 / 12 (50%) of the small EVs (≤ 100 nm) are marked (assuming half of these contain two AuNPs per EV).

**Figure 4:**
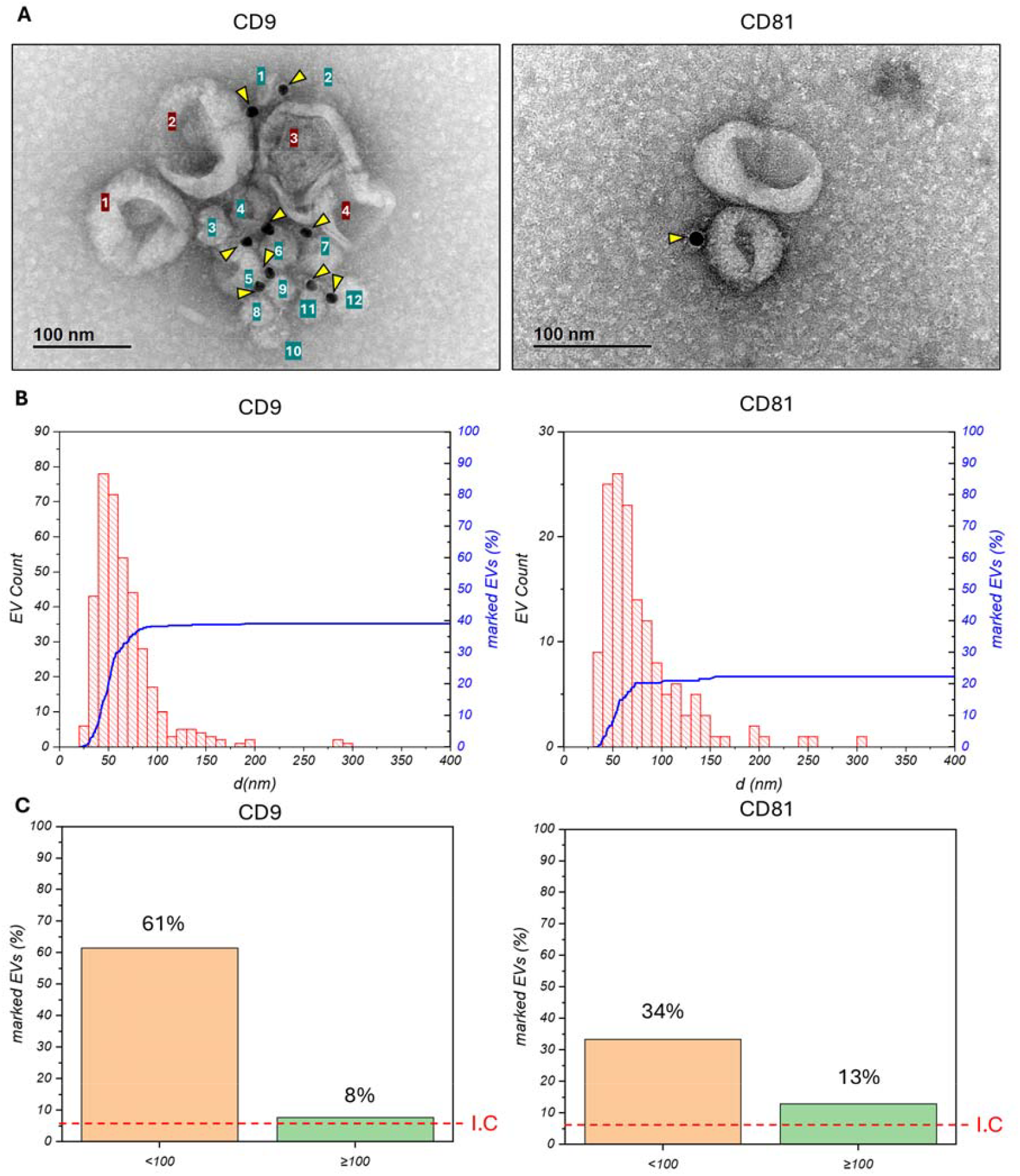
Correlation between EV size and CD9 / CD81 positivity. Representative micrograph of EVs labeled with anti-CD9 (left) or anti-CD81 (right), and treated with AuNP-AG* (A). Yellow triangles indicate bound AuNPs, and numbered labels count small-EVs (green) and large-EVs (brown). The diameter distribution of EVs labeled with each antibody and containing AuNPs on their surface is shown (B), together with the cumulative distribution of the sample (blue line). The percentage of AuNP-labeled EVs is shown in (C), after segregating EVs into two categories based on a diameter cut-off of 100 nm (small EVs: orange; large EVs: green). I.C.: levels in the isotype control (for EVs of any size), representing the background signal of the assay.

To avoid imposing an arbitrary size cut-off to define lEVs and sEVs, we plotted the cumulative distribution of AuNP-positive EVs as a function of the EV diameter. The cumulative positivity for anti-CD9-labeled EVs increased rapidly and stabilized around 90 nm (**Figure 4B)**, not varying significantly for larger sizes. A similar result was obtained for EVs labeled with anti-CD81. Considering that EVs > 100 nm are present in lower proportions, the AuNP-positivity was analyzed by separating populations into two groups, with a cutoff at 100 nm. Results after applying the Z-test for proportions (p=0.05) showed significant differences, both for EVs labeled with anti-CD9 (63% vs 10% for EVs smaller and larger than 100 nm, respectively) and anti-CD81 (35% vs 10%) (**Figure 4C**). These results imply that, while larger EVs do contain CD9 or CD81 above background levels, these tetraspanins are mostly associated with EVs ≤ 100 nm, at least in U-2 OS EVs.

Taken together, our immuno-TEM results explain why the larger EV population could be detected in our preparations by TEM, MRPS, and nanoFCM (**Figure 1A** and **1B**), but not by SP-IRIS (which requires capture using an anti-tetraspanin antibody; **Figure 1C**). More broadly, we present a modified immuno-TEM protocol that does not require the use of commercial gold nanoparticles, and that can be applied to measure the presence and relative abundance of virtually any EV surface protein as a function of EV size.

## 3. Discussion

Conventional immuno-TEM protocols use gold nanoparticles with attached primary antibodies. However, we have found that IgG incubation with AuNP-AG particles synthesized in our lab tend to aggregate during centrifugation, making it difficult to remove excess free antibodies that would otherwise interfere with downstream immunoassays. To circumvent this problem, we incubated EV preparations with primary anti-tetraspanin IgGs, purified the antibody-decorated EVs by SEC, adsorbed these particles into a TEM grid, blocked the grids, and performed immunodetection with BSA-blocked AuNP-AG. In addition, we did not impose any arbitrary size cut-off to our EV preparations by ultracentrifugation (except for an initial clearance step at 2,000 x g). Consequently, we obtained a simple method to determine tetraspanin positivity as a function of EV size in the most unbiased possible manner, using a “ground-truth” technique, TEM. Our protocol could be easily adapted to correlate EV size and the presence of virtually any EV surface marker to which reliable antibodies are available.

Using this modified immuno-TEM protocol, we have shown that CD9 and CD81 are enriched on small-sized EVs (≤ 100 nm), at least for EVs from U-2 OS cells. Positivity rates in small EVs for CD9 and CD81 were 63% and 35%, respectively, compared with only 10% for larger EVs (> 100 nm). These results are consistent with previous studies (Kowal et al., 2016), but our method has the advantage that EVs are analyzed individually in a preparation containing both small and large EVs.

There are several reports using single-vesicle characterization methods to assess variations in surface tetraspanin profiles between EVs from healthy donors and disease patients (Rydland et al., 2023), or as a function of EV source and purification methods (Mizenko et al., 2021). A few reports have analyzed the tetraspanin content of small vs large EVs (Jeppesen et al., 2019; Kowal et al., 2016). However, these reports used EV subpopulations that had previously been separated based on physicochemical parameters such as size and employed bulk analysis methods such as Western blot. Among studies using single-vesicle characterization methods, those based on the ExoView platform with tetraspanin capture chips (Arab et al., 2021; Breitwieser et al., 2022) are unable to see tetraspanin-negative vesicles and, hence, cannot provide results comparable to those shown herein. We have earlier reported that 40% of SEC-purified EVs or ultracentrifugation-purified small EVs (100,000 x g pellet, after prior removal of large EVs at 10,000 x g) were positive for CD81 when sourced from the T lymphocyte line H9 and measured by nanoparticle tracking analysis in fluorescent mode (Arab et al., 2021). This is comparable to the 35% value determined in this work for small EVs (**Table 1**). A study in porcine seminal plasma used flow cytometry to separate small and large EVs (in the paper referred to as exosomes and microvesicles) and then registered the % of particles positive for each tetraspanin (Barranco et al., 2019). Percentages are higher than those found in this study, but the characteristics of the sample and detection method are too different for a direct comparison.

**Table 1.** Information provided by each technique based on its ability to detect EVs according to their size. N.D.: non-determined. N/A: not applicable. (−): non-detected.

|  |  | (immuno)-TEM | nanoFCM | FCM | MRPS | ExoView (SP-IRIS) |
| --- | --- | --- | --- | --- | --- | --- |
| DETECTION | EVs ≤ 100 nm | + | + | - | + | + |
|  | EVs > 100 nm | + | + | + | + | not captured |
| % CD9 | EVs ≤ 100 nm | **63%** | N.D. | 36% | N/A | N/A (ratios only) |
|  | EVs > 100 nm | **10%** |  |  |  | not captured |
|  | All EVs | **40%** |  |  |  | N/A |
| %CD81 | EVs ≤ 100 nm | **35%** | N.D. | - | N/A | N/A (ratios only) |
|  | EVs > 100 nm | **10%** |  |  |  | not captured |
|  | All EVs | **22%** |  |  |  | N/A |

A recent study applied an imaging-based multiplexed assay to SEC-purified EVs derived from different cell lines (Spitzberg et al., 2023). In all cases (PANC-1, CAPAN-2, ASPC-1 and A549 cells), the percentage of EVs positive for CD9 was higher than for CD81, like in this study. For PANC-1-derived EVs, the percentages of CD9+ EVs was 47.9% and the percentage of CD81+ EVs was only 7.3%. While the percentage of CD81+ EVs is lower than in our study (22%), the percentage of CD9+ EVs is comparable to the results shown herein (40%). Another study, using a custom flow cytometer with a limit of sizing as low as 35 nm (Kim et al., 2024), determined the percentage of CD9+ and CD81+ seminal EVs as 34.7% and 4.6%, respectively (Y. Jiang et al., 2021). Interestingly, the authors also analyzed the same samples using super resolution microscopy and arrived at almost identical results. Again, the percentage of CD9 particles is similar to that in our study, but the percentage of EVs positive for CD81 is much lower. Nevertheless, in our study, the ratio between particles captured by CD9 vs. CD81 antibodies is 1.6 based on SP-IRIS results (**Figure S1, D**) and 1.7 based on our immuno-TEM assay (**Table 1**). Thus, discrepancies between our study and results derived from other single-vesicle analysis platforms (Y. Jiang et al., 2021; Spitzberg et al., 2023) are probably explained by differences in the EV source rather than explained by technical biases.

In view of our data, the inconsistent EV size distribution profiles obtained by nanoFCM, MRPS, and SP-IRIS / ExoView can be explained by the inherent limitation of the latter technique as we used it, i.e., with commercially available tetraspanin capture chips. Despite being powerful for multiplex analysis of surface markers at the single-vesicle level, if tetraspanins are the immunocapture target, it cannot capture or detect EVs with low tetraspanin content, which tend to be larger in size. This explains why this technique did not show the bimodal size distribution profile that could be obtained by techniques such as nanoFCM and TEM. Of note, though, the SP-IRIS technique can be adapted to use non-tetraspanin capture methods (Daaboul et al., 2016; L. Jiang et al., 2022; Reddington et al., 2013), but universal capture may remain an obstacle.

Our results also show that, at least in the case of U-2 OS cells, CD9 and CD81 are more associated with small EVs, without taking their biogenic origin into consideration. This reinforces the idea that classification based on size (e.g., small vs. large EVs) is more practical than classification based on biogenesis (e.g, exosomes vs ectosomes), especially when studying biofluids or cell-conditioned medium. Not every paper using the word “exosome” describes the same thing, but “EVs ≤ 100 nm” is non-ambiguous. However, separating EV preparations into two different size categories needs to be justified from either a compositional or a functional perspective. Here, we show that 100 nm is not only an arbitrary round number; it marks the limit between two EV subpopulations characterized by very different tetraspanin content. The extent to which exosomes and ectosomes are enriched in these two subpopulations remains to be addressed.

## 4. Conclusions

In summary, our results suggest that tetraspanins CD9 and CD81 are enriched in small EVs (≤ 100 nm) derived from U-2 OS cells, indicating a correlation between EV size and the presence of these proteins. While this idea has been around for decades (Théry et al., 2002), later work showing the enrichment of CD9 in ectosomes (Mathieu et al., 2021), and the body of literature wrongly describing ectosomes and microvesicles as consistently ≥ 100 nm, motivated us to revisit some of the foundations in the field. We also provide a modified immuno-TEM protocol that can be easily implemented in most labs and can be used to study the correlation between other surface proteins and EV size in different cell types.

## 5. Acknowledgments

The authors would like to thank Paula Céspedes and the Cell Biology Unit of the Pasteur Institute of Montevideo for their help with flow cytometry, and Valentina Blanco for her help with EV purification. PF, JPT, AC and EM are researchers from PEDECIBA (Uruguay) and from the Sistema Nacional de Investigadores (SNI, Uruguay). PF would like to thank Agencia Nacional de Investigación e Innovación (ANII, Uruguay) and Comisión Académica de Posgrado (CAP, UdelaR) for scholarships doing this work.

## 7. Materials and methods

### 7.1 Reagents

For the synthesis of citrate-capped gold nanoparticles (AuNP-cit), chloroauric acid (HAuCl_4_·3H_2_O, 99.9%, Sigma-Aldrich) and sodium citrate (Na_3_C_5_H_6_O_8_, 99%; Carlo Erba) were used. For AuNP modification, bovine serum albumin (BSA, ∼66 kDa, protease-free, > 96%; Capricorn Scientific) and recombinant A/G protein (A/G, ∼50 kDa, 30 mg/mL, Thermo Scientific) were used. For flow cytometry, we used Fc Blocking reagent (Human TruStain FcX™), and the following fluorophore-conjugated antibodies: anti-human CD9 (Biolegend, Cat. No.: 312116, PE/Cy7) and anti-human CD81 (Beckman Coulter B19717, CD81-Pacific Blue, JS64, 0.5 mL, ASR).

For TEM EV assays, mouse anti-human CD9 antibody (IgG1κ, 500 µg/mL, H19a, BioLegend) and mouse anti-human CD81 antibody (IgG2b, 500 µg/mL, MAB4615, R&D Systems) were used. As isotype control, anti-nucleolin antibody (anti-human C23, IgG1k, 200 µg/mL, sc-8031, Santa Cruz Biotechnology) was used.

All protein solutions were prepared in phosphate-buffered saline (PBS, NaH_2_PO_4_/Na_2_HPO_4_ 10 mM, NaCl 150 mM, KCl 2 mM, pH = 7.4). Ultrapure water (18.2MΩ.cm) was used throughout this work.

### 7.2 Synthesis of AuNPs

Synthesis of AuNP-cit was done by the Turkevich method (Turkevich et al., 1951), as in work previously published by our group (Fagúndez et al., 2021). Briefly, 50 mL of ultrapure water and 1 mL of a 20 g/L chloroauric acid solution (approximately 50 μmol) were placed into a round-bottomed two-neck flask. The system was completed with a water condenser and heated to boiling, and 5 mL of a 38.8 mM sodium citrate solution were added immediately. The solution was further heated under reflux until the appearance of an intense red-bordeaux color and was maintained for an additional 10 min. The AuNP solution was allowed to cool in darkness for 24 h before characterization. Characterization of the AuNPs was carried out using transmission electron microscopy (TEM), spectrophotometry (visible range), colloidal stability assay, and electrophoretic mobility to assess the Z-potential, and Dynamic Light Scattering (DLS) to asses the hydrodynamic diameter, as detailed in our previous work (Fagúndez et al., 2021).

### 7.3 Synthesis of AuNP-protein conjugates

AuNP-AG and AuNP-BSA were made by ligand exchange using the adsorption method. One hundred µL of a 200 µg/mL protein solution (in PBS) was added to 1 mL of AuNP-cit suspension and incubated for 24 h at 4°C. Protein concentrations between 10 and 200 µg/mL were tested to ensure nanoparticle surface saturation at the working concentration. The stability of the conjugates was evaluated by UV/Vis spectrophotometry, DLS, colloidal stability assay and electrophoretic mobility according to our previous work (Fagúndez et al., 2021).

### 7.4 Separation of extracellular vesicles (EVs)

EVs were separated from cell-conditioned media (t = 24 h) of human osteosarcoma epithelial cells (U-2 OS). Cells were initially cultured in T75 flasks using DMEM, high glucose with GlutaMax (Gibco) + 10% fetal bovine serum (FBS, Gibco), until reaching 80-90% confluence. Prior to EV collection, the conditioned medium was removed, cells were washed three times with sterile PBS, and 7 mL of serum-free MEGM medium (Lonza; without addition of the bovine pituitary extract and the antibiotics included in the kit) were added. Note that, although this defined medium is commercialized for the growth of mammary epithelial cells, we have used it extensively with U-2 OS cells without evident loss of cell viability or proliferation capacity (Tosar et al., 2020). After 24 h, the cell-conditioned medium was collected and centrifuged at 300 x g to remove detached cells and then at 2,000 x g for 15 min at 4°C to remove cellular debris and very large vesicles. The pellet was discarded, and the supernatant was concentrated by ultrafiltration (Vivaspin 20, Sartorius; 10 kDa, 4°C) to 500 µL. EVs were further purified by size exclusion chromatography (SEC, qEVoriginal, iZON, pore size ≤ 70 nm). Briefly, 500 µL of concentrated supernatant was loaded, and elution was performed by gravity with ice-cold PBS (filtered through 0.2 µm). Columns were pre-chilled by equilibration in cold PBS. A volume of 2.8 mL (void volume) was eluted and discarded, and fractions of 1 mL were collected afterwards. Fractions 1 and 2 (containing the majority of EVs, Arab et al., 2021) were combined and subsequently concentrated using ultrafiltration to 200 µL.

### 7.5 Western blot

The cell-conditioned medium of U-2 OS cells grown in DMEM + 10% FBS was collected, concentrated by ultrafiltration, and underlaid beneath a 12-36% iodixanol density gradient as described earlier (Tosar et al., 2020). The gradients were centrifuged overnight at 120,000 x g and 4°C, using a SW40 Ti rotor. Twelve sequential fractions of 1 mL were obtained, beginning from the top of the gradient. An aliquot of 0.5 mL of each fraction was twice precipitated with cold (−20°C) 60% acetone, and the pellets were resuspended in 1x loading buffer (without reducing agents) for Western blot analysis using the following antibodies: anti-CD9 (Millipore; CBL162; clone MM2/57) and anti-CD81 (R&D; MAB4615; clone 454720).

### 7.6 Microfluidic resistive pulse sensing (MRPS)

To obtain particle number concentrations and size profiles of EV preparations, a resistive pulse sensing device coupled with microfluidics (MRPS, nCS1, Spectradyne) was used. To obtain a representative value of the concentration and size distribution of U-2 OS EVs, a pool (28 mL; 4x T75 flasks) of cell-conditioned media was concentrated down to 2 mL and divided into 4 aliquots of 500 µL, one of which was used for the separation and quantification of EVs by MRPS (the remaining were kept at -20°C for later use). Briefly, after SEC purification (see above), EVs were concentrated by ultrafiltration to 200 µL, and then a 1/10 dilution was prepared in PBS / Tween-20 buffer (0.1% v/v), previously filtered through 0.2 and 0.02 µm the same day of determination. Subsequently, 10 µL of the previous dilution was taken and placed on a pre-calibrated MRPS chip (C400, size range 65-400 nm, Spectradyne), performing the measurement with an integration time of 10 ms, until approximately 3000 events had been collected. Note that low concentrations of Tween in the buffer are needed to improve wettability of the chips and do not affect EV size and concentrations as determined using this instrument (Arab et al., 2021).

For data cleaning, scatter plots were constructed with the parameters diameter (nm) vs. transit-time (ms) based on the signal-to-noise ratio (SN), using OriginPro 2018 software **(Figure S7)**. From these plots, the optimal cutoff values for transit-time and SN parameters were determined, which were subsequently applied for data cleaning and noise elimination in the proprietary software of the equipment (nCS1, Viewer).

### 7.7 Nano flow cytometry (nanoFCM)

The size distribution and concentration of EVs were assessed at JHU using nanoparticle flow cytometry (NanoFCM Co.), using a freshly prepared EV sample purified as described above. Single-photon counting avalanche photodiodes (APD) were used for the detection of side scatter (SSC) of individual particles. The instrument was calibrated for concentration and size using 250 nm silica beads (NanoFCM QC beads) and a Silica Nanosphere Cocktail (NanoFCM, #S16M-Exo), respectively. EV preparations were diluted in PBS, passed through the detector, and events were recorded for 1 minute at a flow rate of 27.5 nL/min. The flow rate and side scatter intensity determined the number and size of corresponding particles, using respective calibration curves. The concentration of EVs was determined from the volume of the analyzed sample.

### 7.8 Characterization of EVs by flow cytometry

Flow cytometry analyses were performed on an Attune NxT cytometer (Thermo Fisher Scientific) with a theoretical nanoparticle diameter limit of detection of 100 nm. Both the blue (488 nm) and violet (405 nm) lasers were used for detection. EVs were separated according to the procedure described above. 200 µL of a freshly prepared EV sample were diluted to 1,000 µL with filtered PBS (0.2 µm) and treated with 5 µL of Fc blocking reagent for 45 min on ice. After this, aliquots of 200 µL were incubated with either 1 µL of PE/Cy7-labeled anti-(h)CD9, or 1 µL of PB-labeled anti-(h)CD81, in the dark. In addition, an aliquot of untreated EVs was analyzed, and the remaining aliquot was diluted with PBS Triton X-100 (final concentration 10% v/v) and incubated for 30 min. The cytometry data were processed with FlowJo software v10.9.0.

### 7.9 SP-IRIS (ExoView)

Characterization of the EV preparations by ExoView was also done at JHU, using an aliquot of the same sample used for nanoFCM, making results directly comparable. The procedure was the same as described by us earlier (Arab et al., 2021). Briefly, an aliquot of SEC-purified EVs was incubated at room temperature for overnight (approx. 16 h) on ExoView chips (NanoView Biosciences, Brighton, MA) that contained the following immobilized antibodies: anti-human CD81 (JS-81), anti-human CD63 (H5C6), anti-human CD9 (HI9a), and, as a negative control, anti-mouse IgG1 (MOPC-21). Subsequently, the chips were incubated for 1 h at room temperature with a cocktail of fluorophore-conjugated antibodies: anti-human CD81 (JS-81, CF555), anti-human CD63 (H5C6, CF647), and anti-human CD9 (HI9a, CF488A) at a dilution of 1:1200 (v/v) in a 1:1 (v/v) mix of incubation buffer and blocking buffer. After several washing stages, the chips were analyzed by interferometric reflectance (SP-IRIS) and fluorescence detection on an ExoView scanner. The data were analyzed using NanoViewer software version 3.1 (NanoView Biosciences).

### 7.10 Characterization of EVs with AuNPs by immuno-TEM

For the immunophenotyping assays, fresh EV samples (obtained within the previous 24 h) were obtained according to the procedure described in section 7.4. For this assay, 500 µL of concentrated supernatant containing EVs were pre-incubated before SEC separation with 1 µL of different antibodies directed against surface tetraspanins, namely: α-CD9 and α-CD81. As an isotype control, EVs incubated with α-nucleolin antibody (1 µL, anti-human C23) were also obtained.

For the immunophenotyping assays, copper grids with carbon film (CF300mesh Cu UL, Electron Microscopy Sciences) were used. First, 5 µL of the marked EV solution was added on top of a UV-treated grid and incubated for 15 min at room temperature. Then the drop was drained with absorbent paper to remove excess liquid. The grid was immediately inverted and placed on the surface of a drop of 0.1% PBS-BSA for 30 min at room temperature to block non-specific sites. The grid was then removed and washed by flotation for 1 min in PBS buffer, drained, and placed on a drop of AuNP-AG or AuNP-BSA solution (1/5 dilution in PBS-BSA, [BSA]f = 20 µg/mL, prepared 30 min before). Subsequently, the grid was washed 3 times (5 min each) with PBS by flotation. After this, it was incubated for 1 min in 2% uranyl acetate solution (negative staining), washed 3 times with ultrapure water, and incubated again in uranyl acetate solution, performing a final wash with ultrapure water. Excess water was removed with absorbent paper and the grid was left to air dry. Observation was carried out on a JEOL JEM 2100 electron microscope at 200 kV (Gatan Orius 1000 CCD camera).

For analysis, an average of 150 images per assay (two assays in total) were taken at a magnification of 80,000x, and they were processed using Digital Micrograph software (Free License, Gatan Microscopy). From each image, a manual count (n = 381 for CD9-labeled EVs using AuNP-AG) of all marked and unmarked EVs was performed, simultaneously recording the number of AuNPs attached to each EV.

Additionally, the size distribution of the EVs was analyzed from the TEM images, measuring the diameter of each EV using the free “measure_features” plugin (Dave Mitchell, http://www.dmscripting.com/measure_features.html). Size distributions were fit to a Normal distribution in OriginPro2018 whenever possible.

