## Supplementary material for "Tetraspanin positivity as a function of extracellular vesicle size measured by a modified immuno-TEM protocol": Figure S1

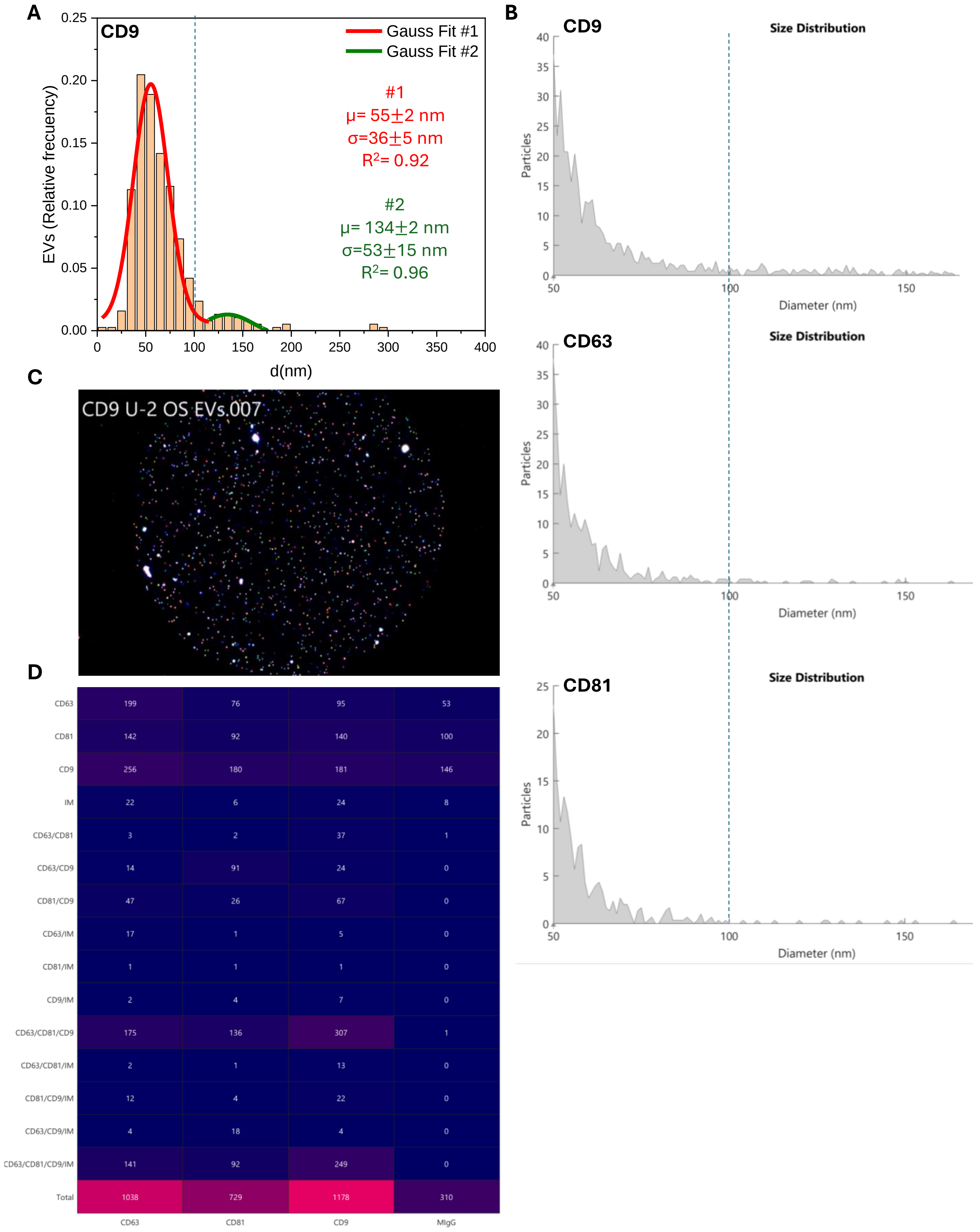


| | CD63 | CD81 | CD9 | MIgG |
|---|---|---|---|---|
| CD63 | 199 | 76 | 95 | 53 |
| CD81 | 142 | 92 | 140 | 100 |
| CD9 | 256 | 180 | 181 | 146 |
| IM | 22 | 6 | 24 | 8 |
| CD63/CD81 | 3 | 2 | 37 | 1 |
| CD63/CD9 | 14 | 91 | 24 | 0 |
| CD81/CD9 | 47 | 26 | 67 | 0 |
| CD63/IM | 17 | 1 | 5 | 0 |
| CD81/IM | 1 | 1 | 1 | 0 |
| CD9/IM | 2 | 4 | 7 | 0 |
| CD63/CD81/CD9 | 175 | 136 | 307 | 1 |
| CD63/CD81/IM | 2 | 1 | 13 | 0 |
| CD81/CD9/IM | 12 | 4 | 22 | 0 |
| CD63/CD9/IM | 4 | 18 | 4 | 0 |
| CD63/CD81/CD9/IM | 141 | 92 | 249 | 0 |
| Total | 1038 | 729 | 1178 | 310 |

**Figure S1: size distribution (TEM, A; SP-IRIS, B) and EV tetraspanin profiling based on the ExoView platform (C, D) .** A) Fitting of the size profile presented in Figure 1A to two Gaussian distributions. B) Size distributions obtained by SPI-IRIS (ExoView platform) using the different capture antibodies. C) Representative image of an anti-CD9 spot in the ExoView platform, incubated with our SEC-purified EVs. Labeled antibodies against CD63 (red), CD81 (green) and CD9 (blue) were incubated with the captured particles. This approach enables to study the co-ocurrance of each tetraspanin on individual particles. The matrix in (D) shows antibody combinations (rows) as a function of the capture antibody (columns). The number of total events in each category is shown.

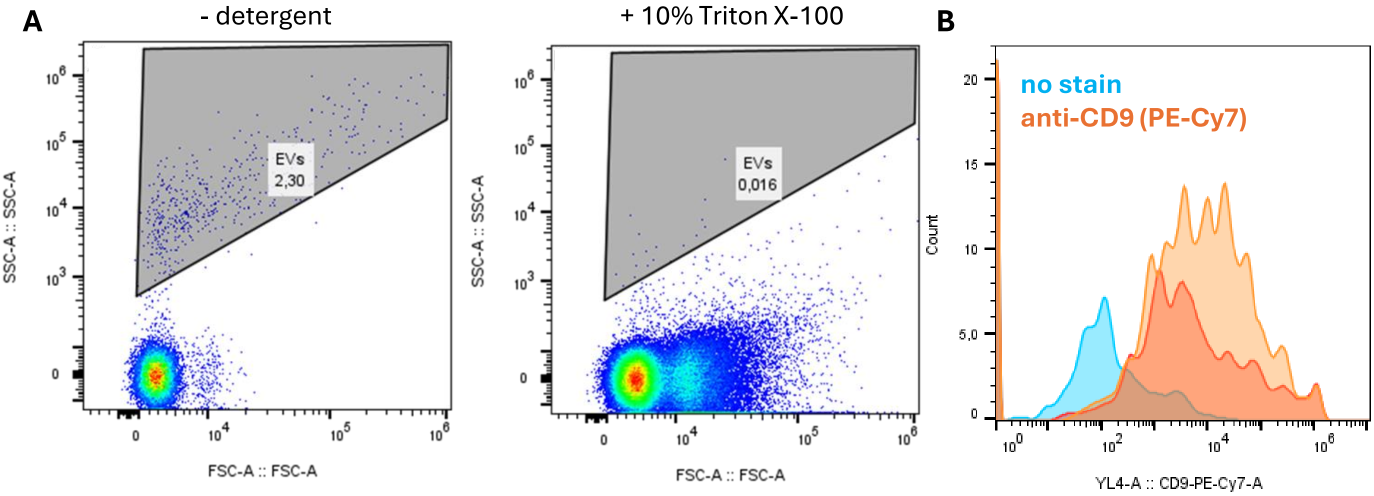


**Figure S2: Flow cytometry analysis (Attune NxT) of SEC-purified EVs.** A) Gating of putative EV events based on the side scatter (SSC) and forward scatter (FSC) using the 488 nm laser. Left: untreated sample. Right: EV preparation treated with. B) Histogram of fluorescence intensity after analyzing gated particles (A) based on the intensity of EV labeled with a PE-Cy7-conjugated anti-(h)CD9 antibody.

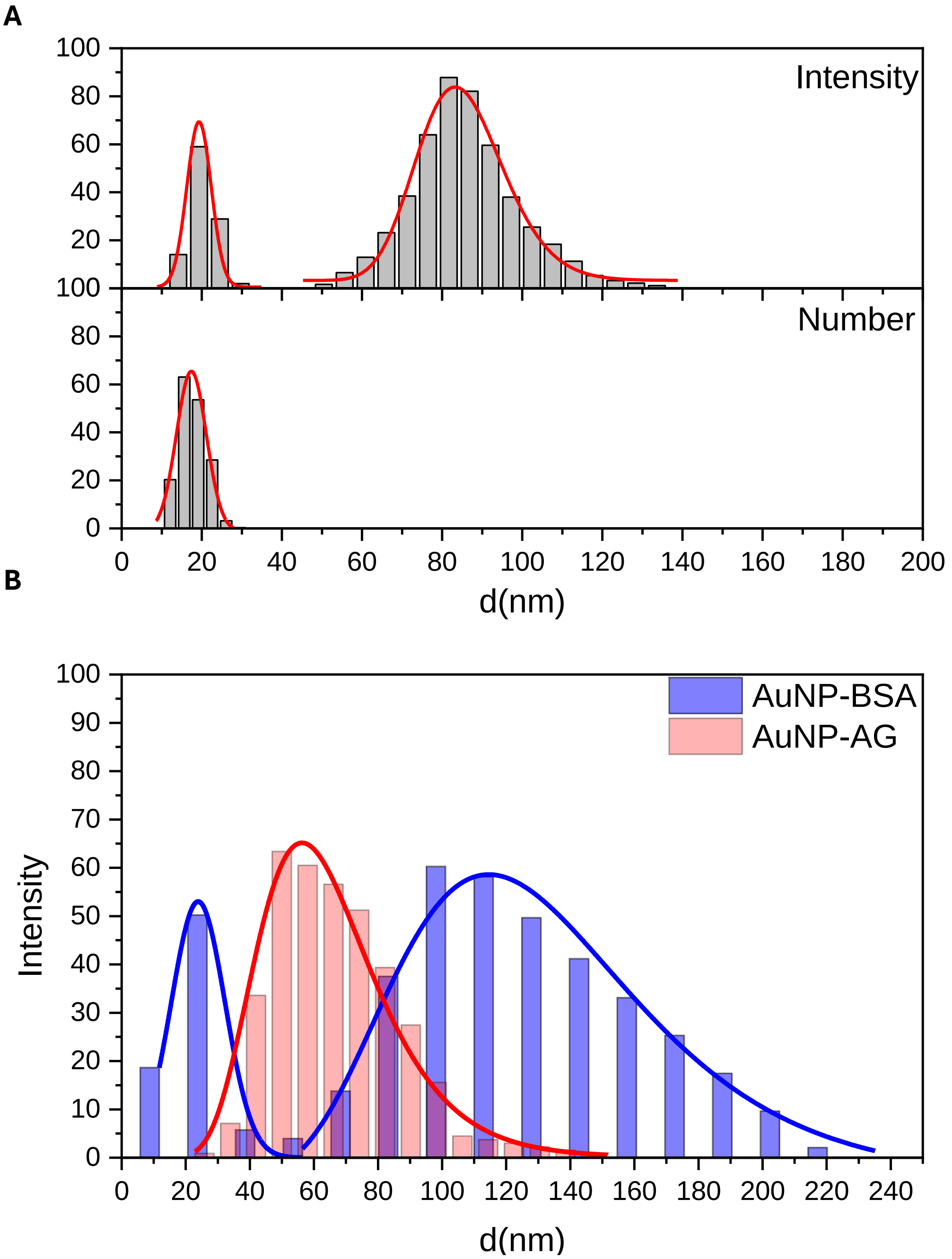


**Figure S3: DLS measurements of unmodified (A) and modified (B) AuNPs.** In (A), the data is presented both by intensity (upper panel) and by number of particles (bottom panel). Note that DLS measurments, when expressed by number, are more representative of the actual size distribution of particles in the sample. Data is presented in (B) in intensity, to show that the AuNP-AG preparation is very stable, with lack of any detectable aggregation products. Data corresponds to AuNPs before centrifugation.

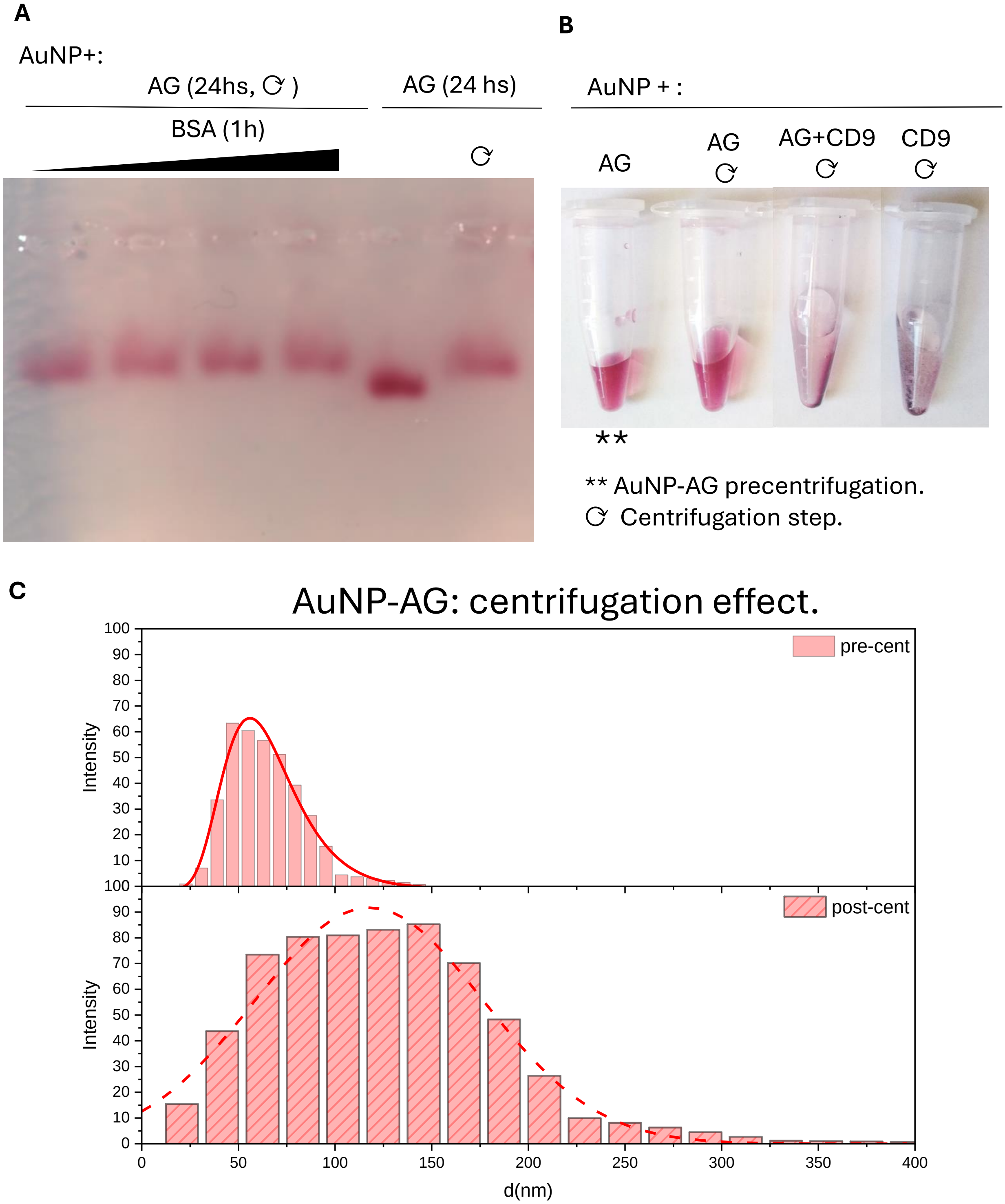


**Figure S4: Agarose gel electrophoresis of protein A/G-modified AuNPs,** either with or without a subsequent BSA blocking step, before and after centrifugation and resuspension in PBS. B) When both AuNP-citrate or AuNP-AG were incubated with an antibody (in this case: an anti-CD9), the particles were no longer stable after centrifugation. C) Even in the absence of an antibody, the size distribution of AuNP-AG is slightly broadened by centrifugation and resuspension in PBS. However, the particles remain red and monodisperse (B, C).

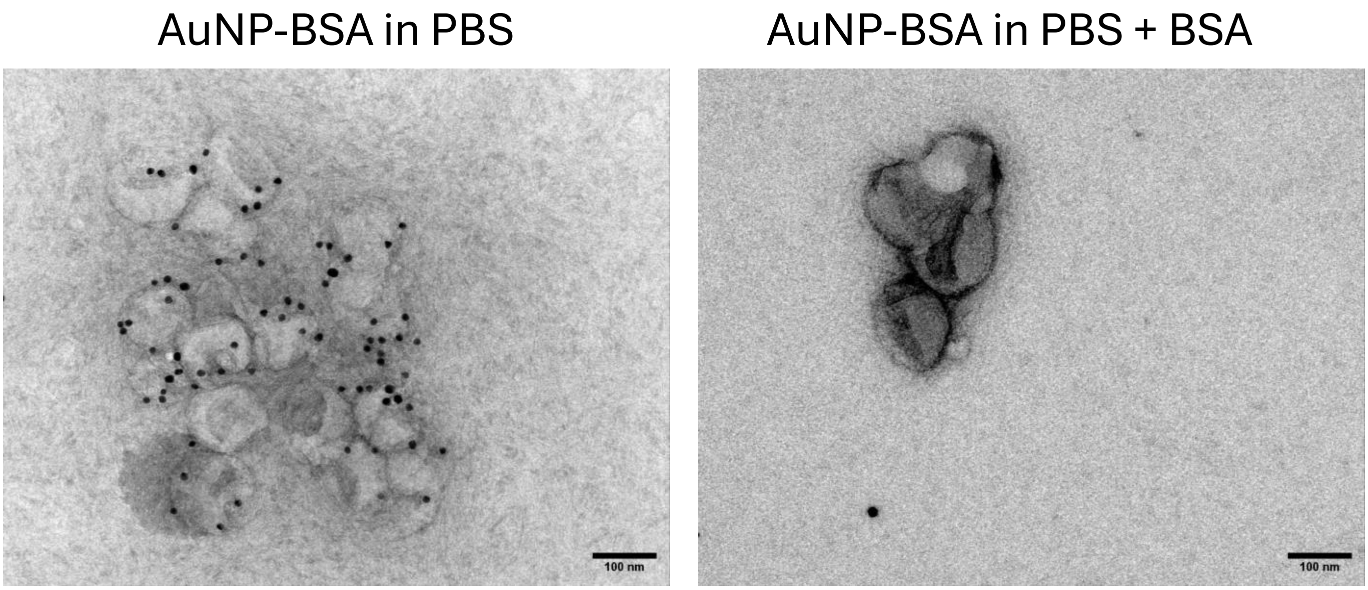


**Figure S5: Incubation of AuNPs in PBS + BSA is essential to prevent unspecific binding.** To study this, we used CD9-labeled EVs and incubated them with AuNP-BSA (i.e., our negative control). Under these conditions, the AuNPs strongly attached to the EVs in an antibody-independent manner (left panel). However, incubating the nanoparticles in the presence of PBS + BSA prevented this undesirable effect (right panel).

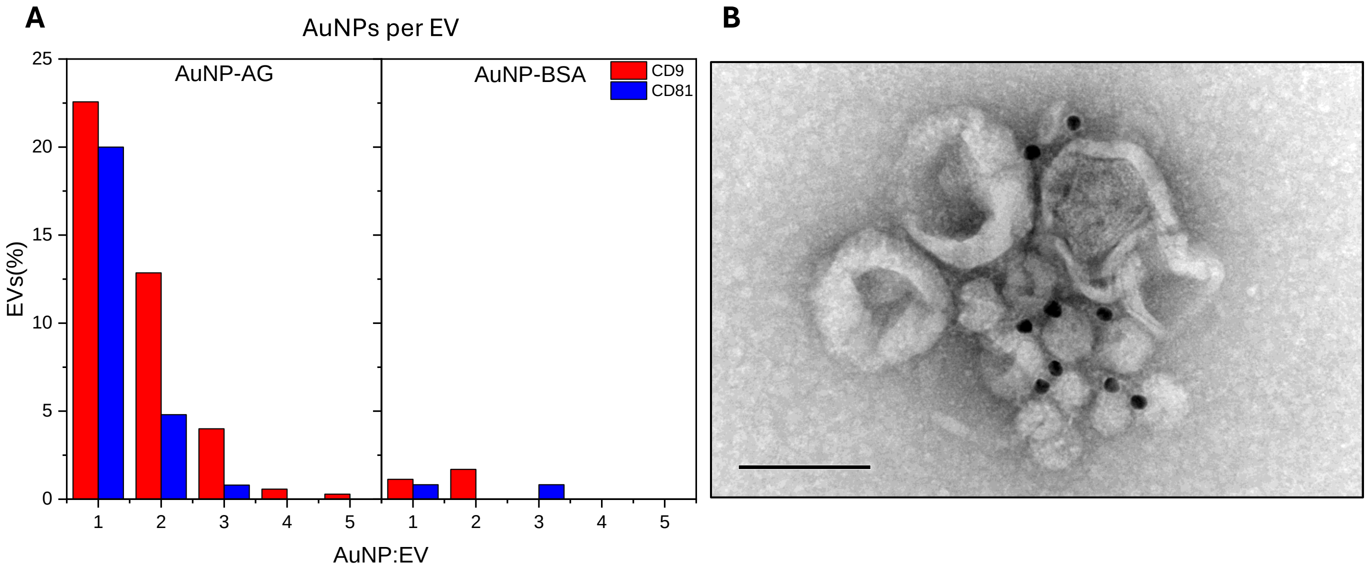


**Figure S6: performance of the modified immuno-TEM method.** A) number of AuNP-AG* (left) or AuNP-BSA (right; negative control) per EV, in CD9-labeled EVs (red) or CD81-labeled EVs (blue). B) Image shown in Figure 4A without labels.

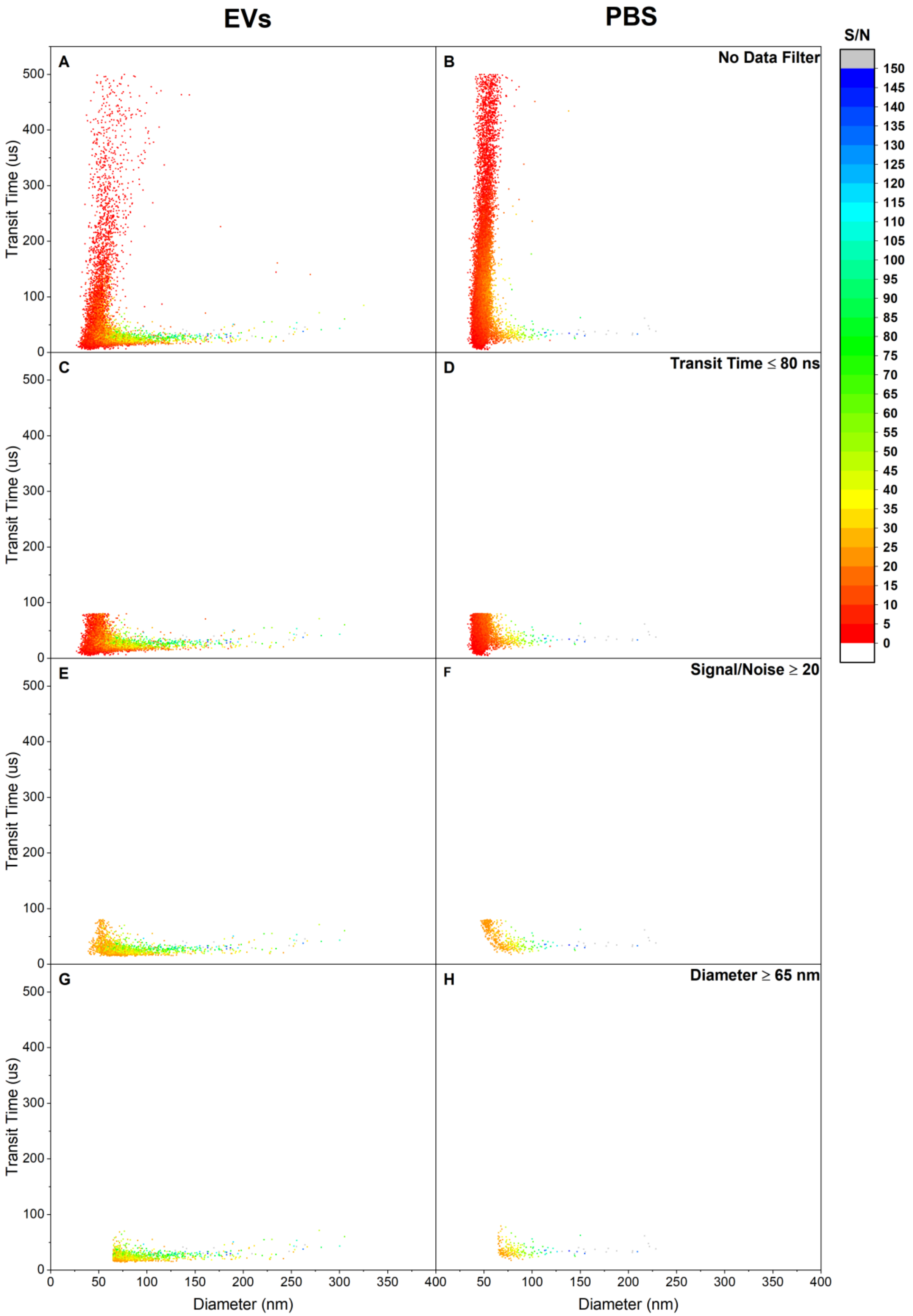


**Figure S7: MRPS results.** Transit time graphs as a function of particle diameter and signal-to-noise ratio. Each panel shows the resulting scatter plot after adding subsequent filters to the data. The measured data from a sample of EVs and a blank composed only of running buffer (PBS-Tween20) are shown for comparative purposes. Filters are (in order): transit time ≤ 80 ns; signal-to-noise ≥ 20; particle diameter ≥ 65 nm.
